# Transcriptome analysis of left versus right intrinsic laryngeal muscles associated with innervation

**DOI:** 10.1101/2023.08.25.554869

**Authors:** Angela M. Kemfack, Ignacio Hernández-Morato, Yalda Moayedi, Michael J. Pitman

**Affiliations:** The Center for Voice and Swallowing, Department of Otolaryngology-Head & Neck Surgery, Columbia University Irving Medical Center, New York, NY; Department of Neurology, Columbia University Irving Medical Center, New York, NY; Pain Research Center, New York University College of Dentistry, New York University, New York, NY

**Keywords:** Laterality, innervation, laryngeal muscles, recurrent laryngeal nerve injury, neuroinflammation, vocal fold paralysis

## Abstract

**Objectives/Hypothesis:** Recurrent laryngeal nerve injury diagnosed as idiopathic or due to short-term surgery-related intubation exhibits a higher incidence of left-sided paralysis. While this is often attributed to nerve length, it is hypothesized there are asymmetric differences in the expression of genes related to neuromuscular function that may impact reinnervation and contribute to this laterality phenomenon. To test this hypothesis, this study analyzes the transcriptome profiles of the intrinsic laryngeal muscles (ILMs), comparing gene expression in the left versus right, with particular attention to genetic pathways associated with neuromuscular function.

**Study Design:** Laboratory experiment.

**Methods:** RNA was extracted from the left and right sides of the rat posterior cricoarytenoid (PCA), lateral thyroarytenoid (LTA), and medial thyroarytenoid (MTA), respectively. After high-throughput RNA-Sequencing (RNA-Seq), 88 samples were organized into 12 datasets according to their age (P15/adult), sex (male/female), and muscle type (PCA/LTA/MTA). A comprehensive bioinformatics analysis was conducted to compare the left-right ILMs across different conditions.

**Results:** 774 differentially expressed genes (DEGs) were identified across the 12 experimental groups, revealing age, sex, and muscle-specific differences between the left versus right ILMs. Enrichment analysis of Gene Ontology (GO) and Kyoto Encyclopedia of Genes and Genomes (KEGG) pathways implicated several genes with a left-right laryngeal muscle asymmetry. These genes are associated with neuronal and muscular physiology, immune/inflammatory response, and hormone control.

**Conclusion:** Bioinformatics analysis confirmed divergent transcriptome profiles between the left-right ILMs. This preliminary study identifies putative gene targets that will characterize ILM laterality.

**Level of Evidence:** N/A

**LAY SUMMARY:** Vocal fold paralysis is more common on the left. This study shows left versus right differences in gene expression related to innervation, suggesting the increased rate of left recurrent laryngeal nerve paralysis may be associated with genetic differences, not just nerve length.

## INTRODUCTION

Normal vocal fold function is necessary for the fundamental activities of daily life: eating, breathing, and speaking. Recurrent laryngeal nerve (RLN) injury results in vocal fold paralysis due to synkinetic reinnervation^[1]^. Vocal fold paralysis causes dysphagia and dysphonia, which leads to frustration, isolation, fear, lost worker productivity, and altered self-identity^[2-5]^. While current treatments have adequate results, they are suboptimal and disability claims are at rates similar to those of chronic diseases like congestive heart failure, chronic obstructive pulmonary disease, and severe depression^[2, 3]^.

Efforts to restore normal vocal fold function focus on the guidance of reinnervating injured axons back to the correct laryngeal muscle to prevent synkinetic reinnervation and the resultant paralysis^[1, 6]^. Understanding the genes expressed by ILMs related to innervation and reinnervation in the healthy state and after RLN injury may identify putative molecular pathways involved in guiding innervation. These genes can be investigated as therapeutic targets for manipulation after nerve injury to guide axons to the appropriate muscle and restore normal vocal fold function.

Robust clinical evidence supports a greater incidence of left vocal fold paralysis versus right ^[7-15]^. Sidedness of the paralysis has been hypothesized to result from the longer length of the left RLN compared to right. Further, the left RLNs course through the thorax may also increase the risk of injury from thoracic disease and surgery compared to the right RLN^[16]^. Left RLN reinnervation is more demanding than on the right side because it requires greater energy expenditure for protein translation and transport from the cell body to the nerve growth cone ^[17-20]^. Intra-axonal transport also implicates the reliance on rapid mitochondrial transport to areas along the axon to meet high energy demand^[21, 22]^. This phenomenon would make the longer left RLN harder to regenerate after acute injury compared to the right.

Cases of idiopathic vocal fold paralysis and paralysis after short-term intubation for surgery also show similar discrepancies in incidence between sides^[23-26]^. These are states where the RLN extension and trajectory should not impact the rate of injury. This persistent sidedness suggests intrinsic neuromuscular differences between the left and right sides may also play a role in the frequency of paralysis^[26]^. In addition, previous findings indicate that intrinsic laryngeal muscles (ILM) exhibit different fiber-type compositions on the left versus the right side in equines^[10]^. Considering that targeted axon outgrowth to the ILMs requires a dialogue between the motoneurons and muscle fibers, an analysis of ILM gene expression may provide insights into the asymmetry of the larynx. These changes in gene expression could also explain the sidedness of vocal fold paralysis^[27]^. Moreover, organ asymmetry has already been identified in the diaphragm muscles and the phrenic nerves that innervate them^[28]^. If intrinsic neuromuscular differences in the larynx (aside from length) influence disparities in reinnervation between sides, these differences may be exploited as therapeutic targets.

The present work aims to investigate inherent neuromuscular differences by testing the hypothesis that, in uninjured laryngeal muscles, there are asymmetric discrepancies in gene expression related to neuromuscular function. Such genes could contribute to the disparity in the incidence of left versus right vocal fold paralysis and warrant further investigation. This study uses recently described techniques to evaluate differences in the transcriptome between the respective left-right posterior cricoarytenoid (PCA), lateral (LTA), and medial thyroarytenoid (MTA) muscles in an uninjured rat model to establish the baseline of gene expression in the healthy animal^[29]^.

## MATERIALS AND METHODS

### Rats and Housing

A total of 82 Sprague Dawley rats were used in this study and distributed in 12 groups according to age (adult and P15), sex (male and female), and type of muscle (PCA, LTA, and MTA). A schematic representation of the experimental design is illustrated in Figure 1. This study was performed in compliance with the Public Health Service Policy on Humane Care and Use of Laboratory Animals, the National Institutes of Health Guide for the Care and Use of Laboratory Animals, the Animal Welfare Act (7 U.S.C. et seq.), and the ARRIVE guidelines. The Institutional Animal Care and Use Committee of Columbia University Medical Center approved the animal use protocol.

**Figure 1.**
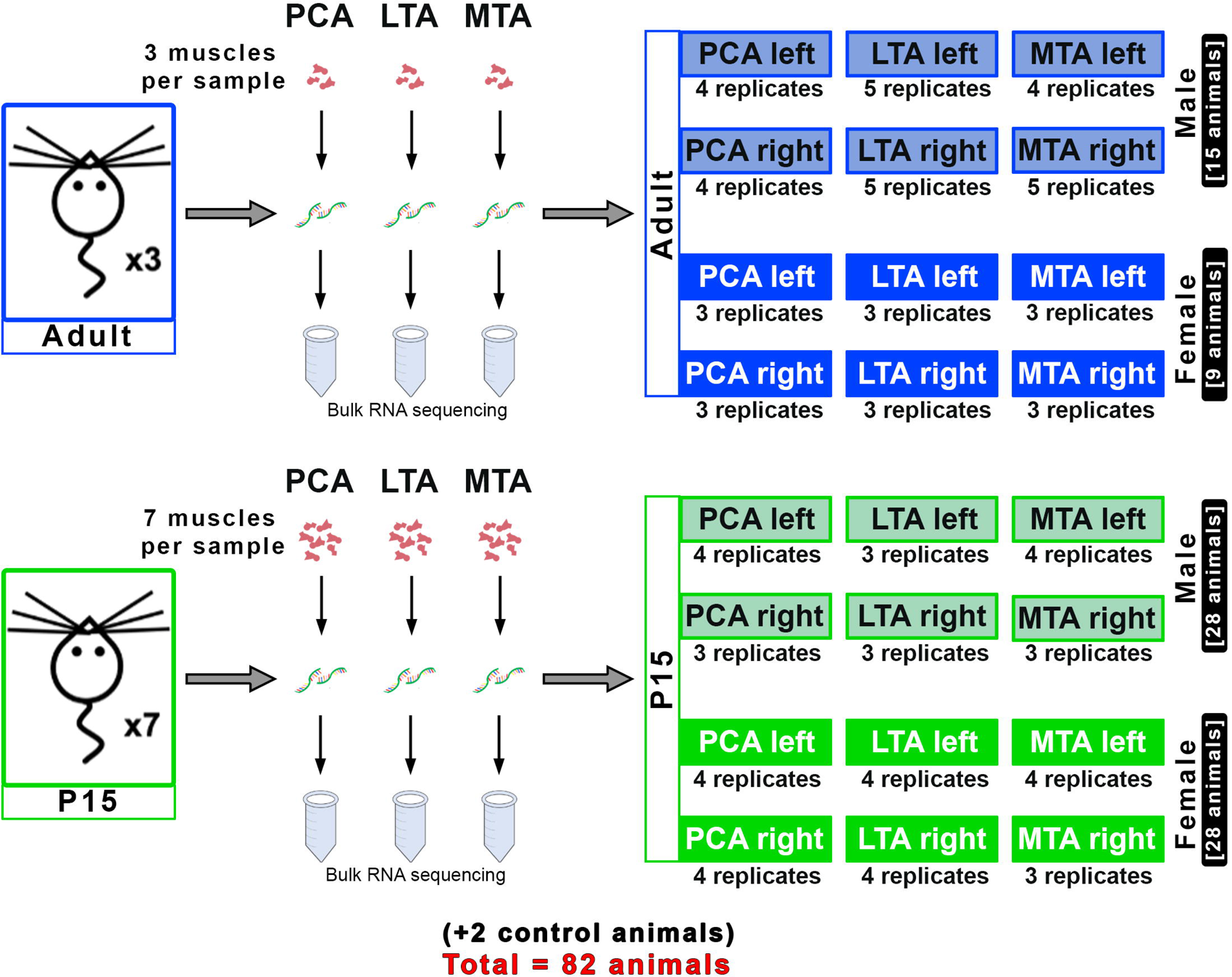
A schematic diagram illustrating the experimental design for high-throughput RNA-Sequencing (RNA-Seq) of the distinctive intrinsic laryngeal muscles.

### Surgical Procedure and Tissue Processing

All rats were euthanized via an overdose of ketamine and xylazine before the larynx was dissected out. ILM isolation for RNA-Seq was performed as previously described [29]. Harvested larynxes were stabilized by briefly rinsing in 2 mL of RNA*later* (catalog #AM7021, Invitrogen, Waltham, MA, U.S.A.) prior to dissection of the muscle bellies using a Leica S8 APO compound microscope (Leica Microsystems, Buffalo Grove, IL). With micro-forceps and micro-scissors, the left and right PCA, LTA, and MTA were carefully excised and stored overnight at 4 C° in 1 mL RNA*later*. The next day, excess stabilizing reagent was removed and the muscle bellies were placed into BioMasher III (Nippi, Inc., Japan) homogenization tubes according to the age, sex, and muscle type (Figure 1). Samples were physically disrupted in liquid nitrogen before adding lysis buffer from the Aurum Total RNA Mini Kit (catalog #732-6820, Bio-Rad, Hercules, CA, U.S.A.). Further homogenization was conducted using a TissueRuptor II (catalog #9002755, Qiagen, MD). All proceeding steps were according to the commercial RNA isolation protocol. Total RNA was measured using a micro-volume spectrophotometer NanoDrop ONE (catalog #ND-ONE-W, Thermofisher, Wilmington, DE). Agilent RNA 6000 Pico Kit (Eukaryote Total RNA Pico, Agilent Technologies, Santa Clara, CA, U.S.A.) and Agilent 2100 bioanalyzer (Agilent Technologies, Santa Clara, CA, U.S.A.) were used to evaluate the RIN. The recovered RNA used in the subsequent sequencing steps reached a RIN value of 7.9 _±_ 0.4 (*N* = 90).

### Library Preparation and RNA Sequencing

The following RNA-Sequencing procedures were performed using the Illumina NextSeq 500 (Illumina Inc., San Diego, CA, U.S.A.) desktop sequencer at the Columbia University Genome Center. Standard analytics were obtained by first using a poly-A pull-down method to enrich mRNA from the total RNA within the sample. Library construction then proceeded using TruSeq chemistry and was sequenced by the NovaSeq 6000 platform (Illumina, Inc). Samples were multiplexed in each lane and pseudo-aligned to a kallisto (0.44.0) index created from human (GRCh38) and mouse (GRCm38) transcriptomes. After quality control steps, the pseudo-aligned reads were mapped to the annotated *Rattus norvegicus* genome assembly Ensembl database.

### Measured Expression Analysis

Differential gene expression analysis in the left versus the right respective ILM was performed through the EdgeR Bioconductor Package 3.32. Sample counts were normalized with qsmooth. Limma-zoom was used to evaluate the sample weights and downweigh the effect of sample outliers. A comprehensive list of differentially expressed genes (DEGs) was generated under three different conditions in the left versus the right comparison. 88 samples, not including the controls, were dichotomized into “left” vs. “right” groups according to the muscle, age, and gender of the rat. The total number of DEGs exhibited using False Discovery Rate (FDR)<0.05 and log fold change >|0.6| across all rat cohorts is displayed in **Table 1**. Raw data can be found in the Gene Expression Omnibus (https://www.ncbi.nlm.nih.gov/geo/query/acc.cgi?acc=GSM6703793).

### iPathwayGuide Analysis

Advaita Bioinformatics iPathwayGuide analysis identified significant genes based on a log fold change of expression with an absolute value of at least 0.6 and a 0.05 threshold for the p-value. Data were analyzed in the context of pathways obtained from the Kyoto Encyclopedia of Genes and Genomes (KEGG) database (Release 100.0+/11-12, Nov 21). Gene Ontologies were obtained from the Gene Ontology Consortium Database. Impact analyses for pathway analysis were evaluated by over-representation of differentially expressed genes in a given pathway and the perturbation of that pathway computed by propagating the measured expression changes across the pathway topology. Pathways from the P15 and adult datasets were compared to identified systematic commonalities.

## RESULTS

Sequencing of the transcriptome across the ninety ILM samples resulted in a mean of 32.9 million mapped reads and a 79.1% mean mapping ratio, with no less than 70% sequence alignment in any sample. Principal component (PC) analysis by experimental condition (i.e., sex, age, ILM) indicated that PC1 and PC2 accounted for 98% variance. Samples were normalized, then, using a mean-variance trend into precision weights, the effects of sample outliers were downweighed. Further quality control measures validated the differential control analysis (Figure 2A).

**Figure 2.**
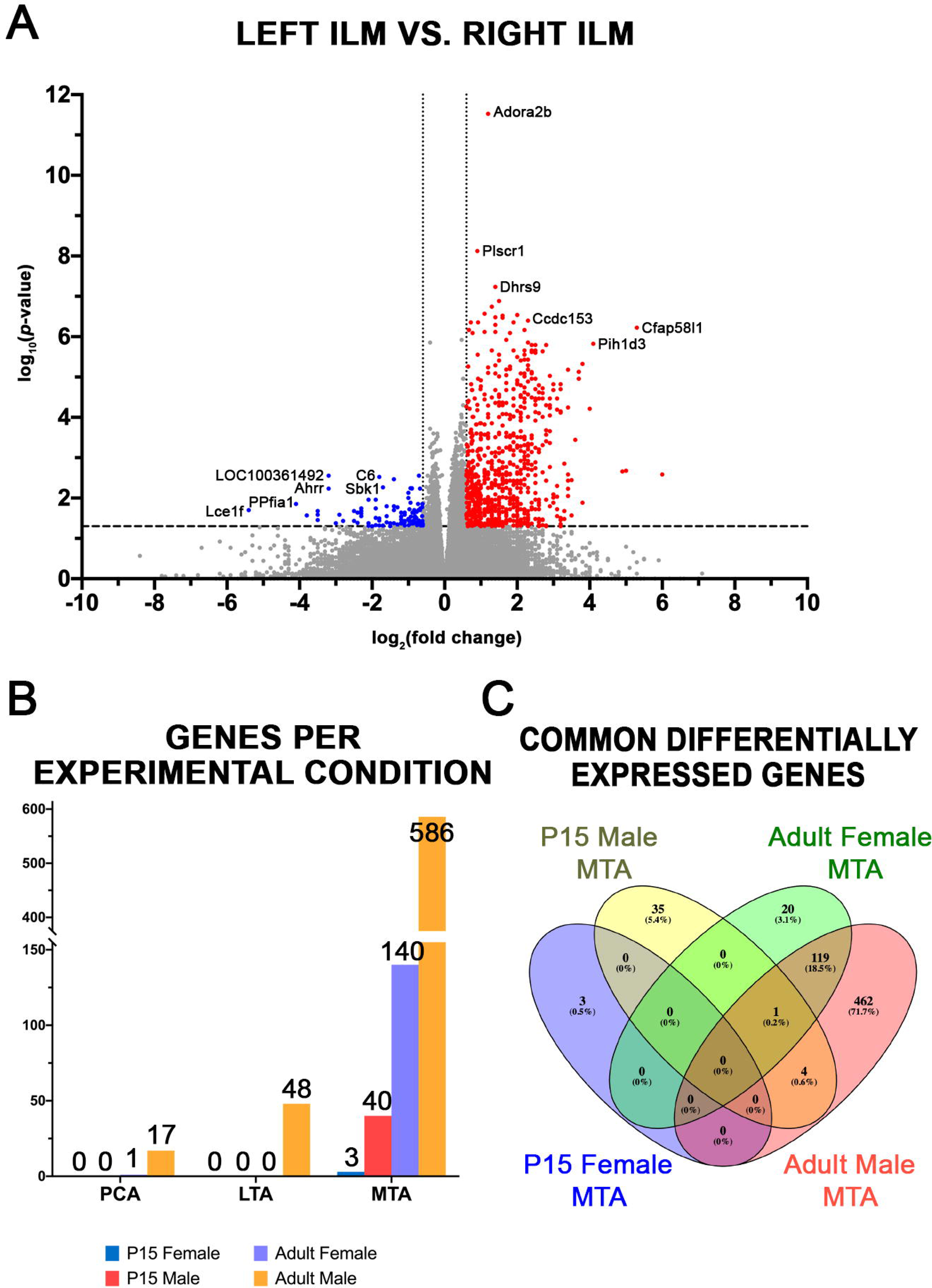
Global differential expression analysis summary comparing gene regulation in the left versus right ILMs. (A) Volcano Plot showing statistically significant genes and their fold change in gene expression across the P15 and adult PCA, LTA, and MTA muscles in both male and female rats. Downregulated differentially expressed genes (DEGs) on the left compared to the right ILMs are represented in blue, while upregulated DEGs are depicted in red. DEGs that were not significant (p-value > 0.05; log fold change > |0.6|) are shown in gray. (B) Number of DEGs in the left compared to the right across each experimental condition. (C) Venn diagram indicating the number of genes that overlapped across all MTA datasets. Common DEGs across the global data intersected three datasets in the MTA muscle.

### Overview of Differential Gene Analysis

Differential expression analysis confirmed asymmetric gene transcriptome profiles between the left-right ILMs according to age, sex, and muscle. Across all PCA and LTA experimental groups, DEGs showed only lower expression (downregulation) on the left versus the right. In contrast, at least 63% of DEGs in the P15 and 95% in the adult MTA were higher (upregulated) on the left relative to the right side. Overall, adult males exhibited the most substantial number of DEGs in each muscle type, with comparatively few DEGs identified in adult females or at P15 in either sex (Figure 2B). Excluding the MTA, which showed overlapping significant genes, DEGs varied per experimental condition (Figure 2C). Only the adult male experimental groups showed overlapping KEGG pathways (Figure 3-6). Common KEGG pathways were only identified Generally, the function of the DEGs per each experimental condition fell under two broad categories: neurophysiological and myogenic factors (Figure 3-6). Gene Ontology (GO) and KEGG Pathways also evidenced asymmetric differences related to inflammatory/immune response and hormonal control. These findings varied according to age, sex, and distinctive ILM.

**Figure 3.**
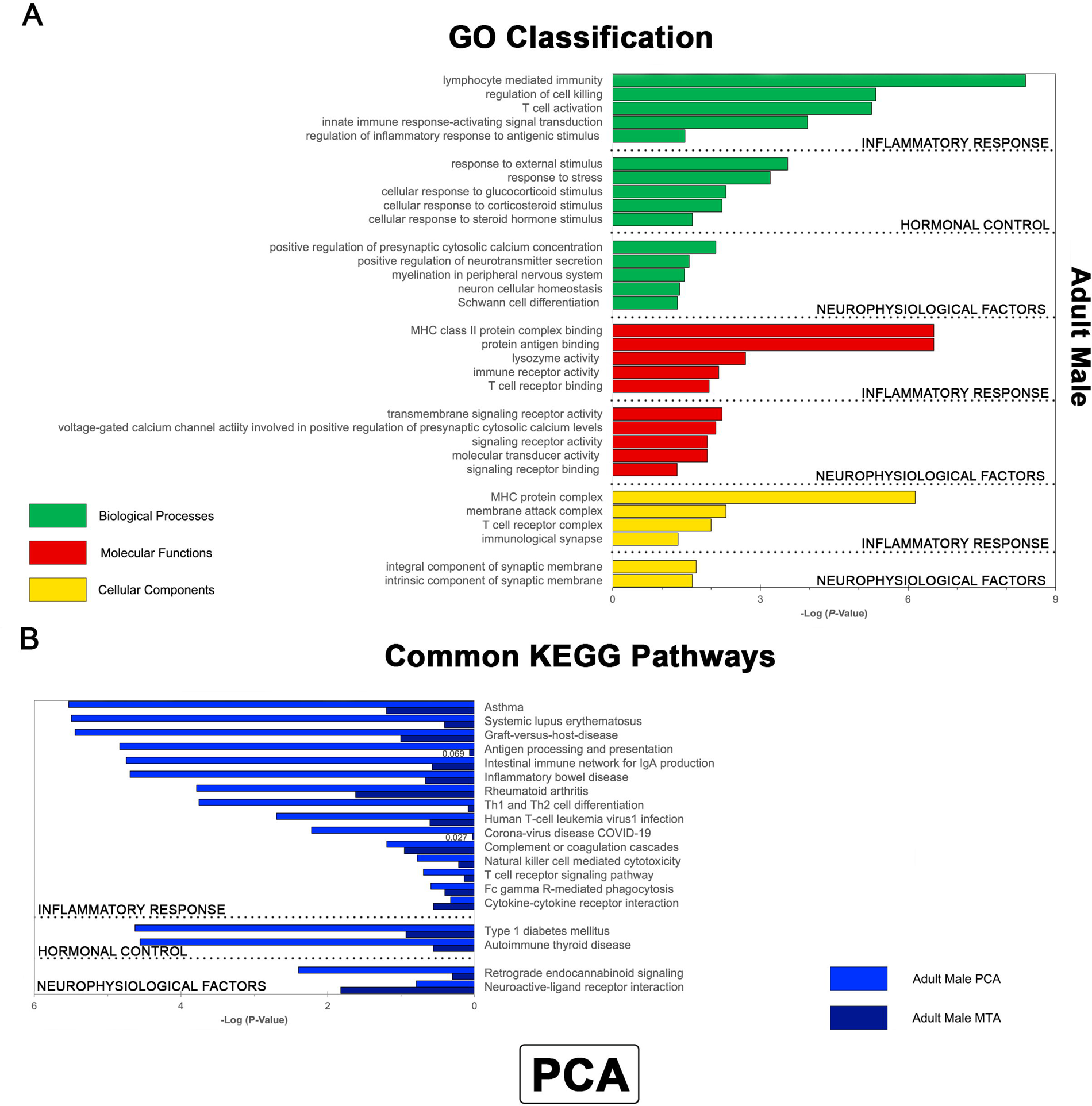
Key Gene Ontology and KEGG pathways enriched in the PCA muscle. (A) Top Gene Ontologies (GOs) enriched in the adult male left vs. right PCA muscle are classified by inflammatory, hormonal, and neurophysiological sub-types in order of statistical significance. Biological processes are shown in green bars; molecular functions are represented in red bars, and cellular components are depicted in yellow bars. (B) Common KEGG pathways enriched in the left-right PCA were only identified between the adult male PCA and adult male MTA datasets. Inflammatory, hormonal, and neurophysiological sub-types are represented by statistical significance. Biological pathways enriched in the adult male PCA dataset are shown as blue bars and those of the adult male MTA dataset are depicted by dark blue bars.

#### PCA

Across the global data, the PCA showed the fewest DEGs of any comparison (Figure 2B). GO Classifications and KEGG Pathways in the left-right adult male PCA indicated that DEGs were highly associated with inflammatory/immune response mechanisms, hormonal control, and neurophysiological processes (Figure 3). Common KEGG pathways between the adult male PCA and MTA datasets showed considerable enrichment of DEGs involved in inflammatory/ immune signaling (Figure 3B). The top pathways of the adult male included ‘antigen processing and presentation’ in addition to ‘Th1 and Th2 cell differentiation’, which are established inflammatory/immune response mechanisms. Interestingly, differential expression analysis in the left-right adult male PCA did not implicate the involvement of myogenic processes in the PCA asymmetry. The adult female PCA only exhibited differential expression of a single mitochondrial iron ion transporter gene, Slc25a37, between sides and thus was not amenable to further GO or biological pathway analysis.

#### LTA

The adult male LTA dataset was the only condition to exhibit significant differential gene expression in the left compared to the right. GO and KEGG pathway enrichment analysis indicated that DEGs represented in the adult male LTA were highly associated with neurophysiological, myogenic, inflammatory, and hormonal regulation (Figure 4). Specifically, top GOs included the ‘semaphorin-plexin signaling pathway’, a hallmark of axon guidance, and the ‘postsynaptic neurotransmitter receptor cycle’ (Figure 4A). Top myogenic factors and inflammatory response included ‘satellite cell activation involved in skeletal muscle regeneration’ and ‘negative regulation of necrotic cell death’, respectively (Figure 4A). Top KEGG Pathways demonstrated highly associated DEGs involved in inflammatory/immune response, such as the ‘IL-17 signaling pathway’, and neurophysiological processes, such as the ‘calcium signaling pathway’ (Figure 4B).

**Figure 4.**
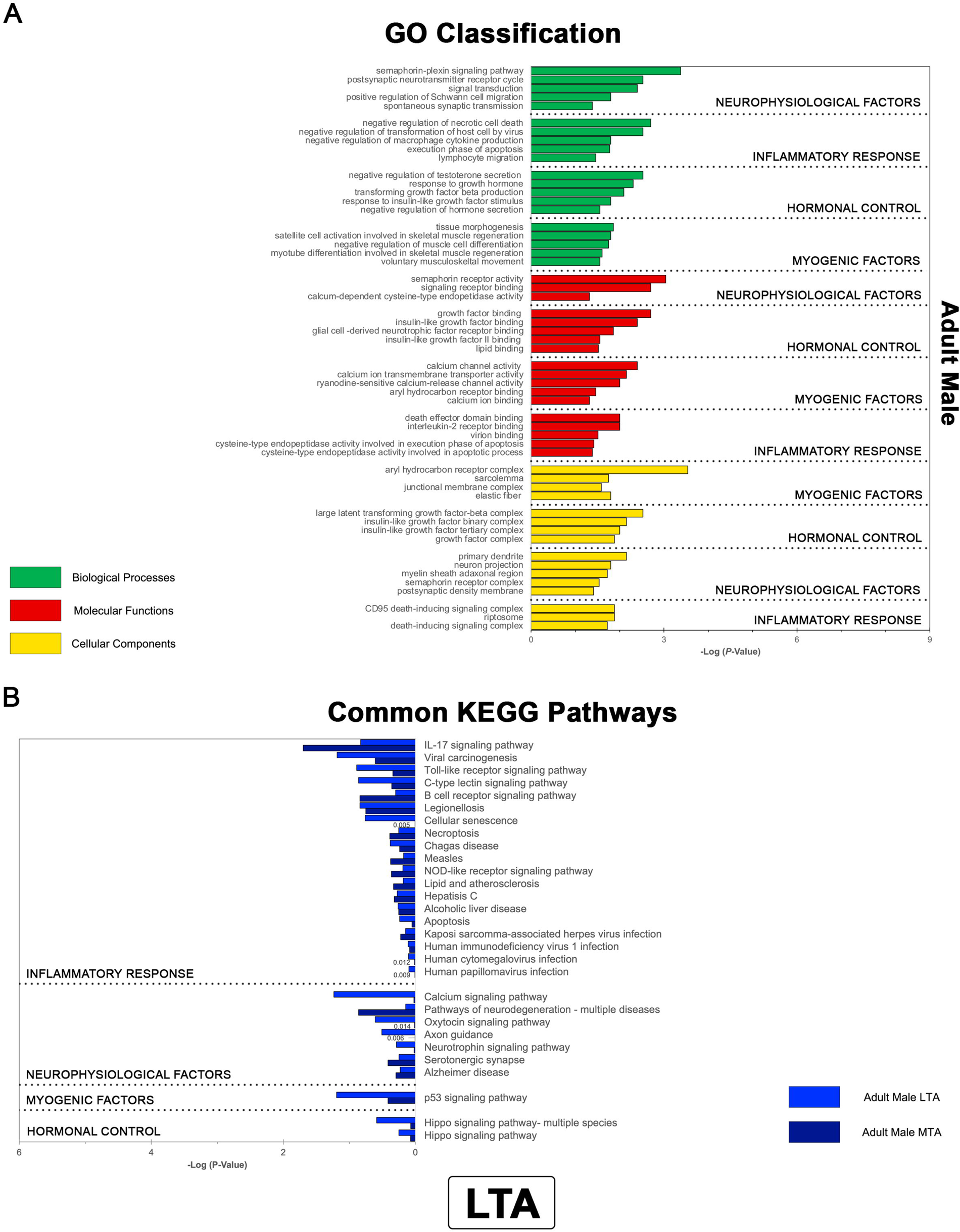
Key Gene Ontology and KEGG pathways enriched in the LTA muscle. (A) Top Gene Ontologies (GOs) enriched in the adult male left vs. right LTA muscle are classified by neurophysiological, inflammatory, hormonal, and myogenic sub-types in order of statistical significance. Biological processes are shown in green bars; molecular functions are represented in red bars, and cellular components are depicted in yellow bars. (B) Common KEGG pathways enriched in the left-right LTA were only identified between the adult male LTA and the adult male MTA datasets. Inflammatory, neurophysiological, myogenic, and hormonal sub-types are represented by statistical significance. Biological pathways enriched in the adult male LTA dataset are shown as blue bars and those of the adult male MTA dataset are depicted by dark blue bars.

#### MTA

Overall, the MTA showed the most DEGs of any condition (Figure 2B). DEGs in this muscle group were identified across sexes in both the adult and P15 datasets, although the high number of DEGs intersecting between the adult male and female MTA is most likely attributable to residual mucosa surrounding the muscle during dissection. GO Classification in the P15 and adult male and female MTA datasets demonstrated the enrichment of inflammatory, neurophysiological, myogenic, and hormonal processes (Figure 5A-D). However, no common pathways that regulate hormonal control were found between the adult male and female datasets (Figure 5E). Similar KEGG Pathways that direct hormonal control were identified between the adult and P15 male MTA (Figure 5D, E). Top GO Classifications in the P15 male MTA included relevant myogenic and neurophysiological factors, such as ‘regulation of muscle adaptation’ and ‘inhibitory chemical synaptic transmission in response to denervation’. Relevant neurological pathways across the adult male, adult female, and P15 male MTA included ‘neuroactive-ligand receptor interaction’ and ‘adrenergic signaling in cardiomyocytes’, which may be similar to pathways in striated muscles that are less characterized (Figure 5C).

**Figure 5.**
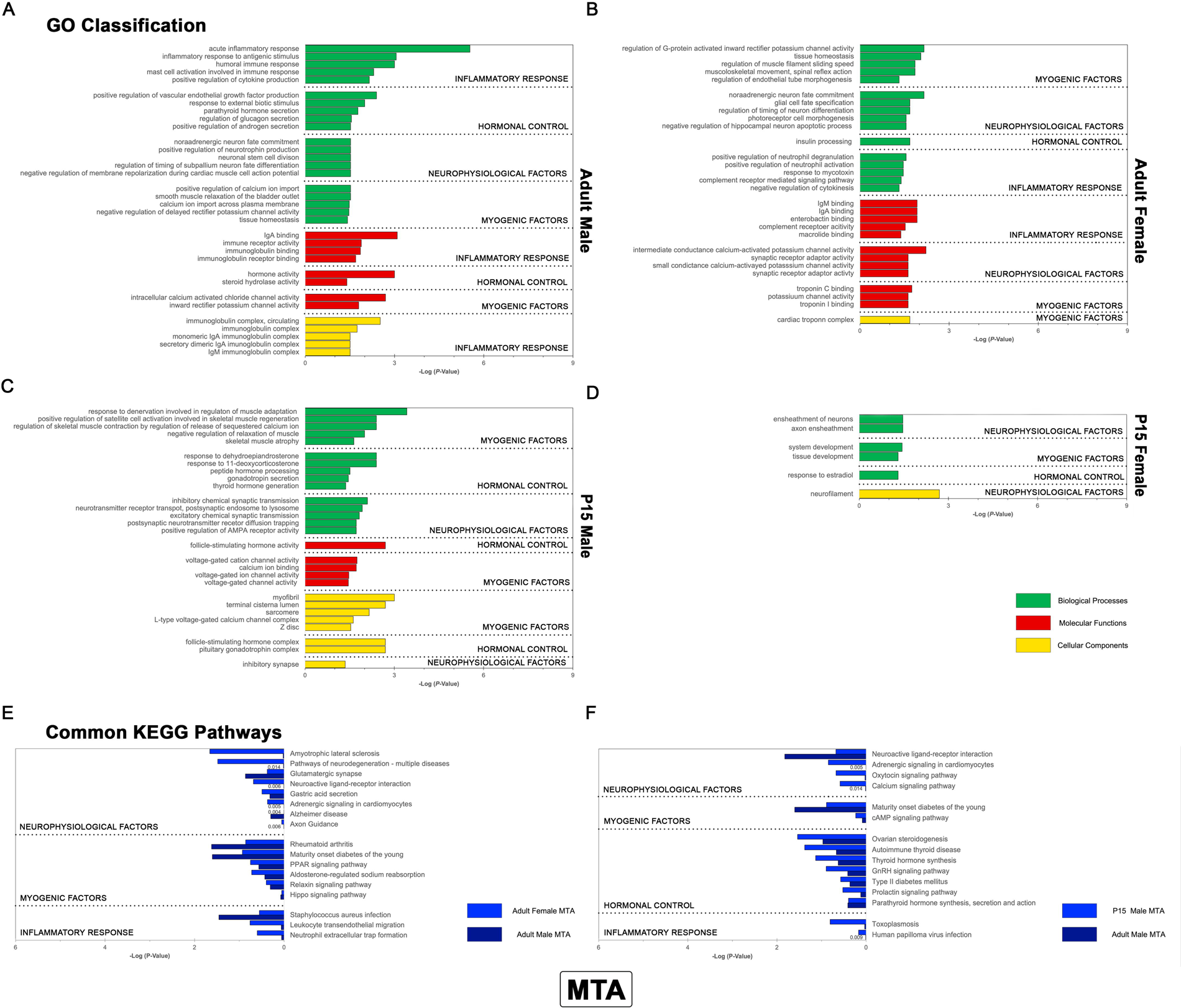
Key Gene Ontology and KEGG pathways enriched in the MTA muscle. (A) Top Gene Ontologies (GOs) enriched in the adult male left vs. right MTA are classified by inflammatory, hormonal, neurophysiological, and myogenic factors in order of statistical significance. (B) Top Gene Ontologies (GOs) enriched in the adult female left vs. right MTA are classified by myogenic, neurophysiological, hormonal, and inflammatory factors in order of statistical significance. (C) Top Gene Ontologies (GOs) enriched in the P15 male left vs. right MTA muscle are classified by myogenic, hormonal, and neurophysiological sub-types in order of statistical significance. (D) Top Gene Ontologies (GOs) enriched in the P15 female left vs. right MTA muscle are classified by myogenic, hormonal, and neurophysiological sub-types in order of statistical significance. (A-D) Biological processes are shown in green bars; molecular functions are represented in red bars, and cellular components are depicted in yellow bars. (E-F) Common KEGG pathways enriched in the left-right adult male MTA intersected all other adult male datasets in addition to the adult female MTA and P15 male datasets. (E) Neurophysiological, myogenic, and inflammatory sub-types are represented by statistical significance. Biological pathways enriched in the adult female MTA dataset are shown as blue bars and those of the adult male MTA dataset are depicted by dark blue bars. (F) Neurophysiological, myogenic, hormonal, and inflammatory sub-types are represented by statistical significance. Biological pathways enriched in the P15 male MTA dataset are shown as blue bars and those of the adult male MTA dataset are depicted by dark blue bars.

#### Age-related differences

P15 muscles generally demonstrated less significant left-right differences in gene expression compared to the adult conditions. The PCA and LTA showed no lateral differences at P15 (Figure 2B). Markers of inflammatory regulation showed marginal differential expression in P15 rats compared to adults (Figure 5C, D, F). The involvement of immune response mechanisms was demonstrated in every adult experimental condition (Figure 3B, 4B, 5E).

#### Sex-specific changes

Left-right differences corresponding to sex were primarily distinguished in the adult male MTA, given the lack of significant DEGs in the female PCA and LTA muscles. Differential expression analysis consistently exhibited enriched pathways and gene ontologies involved in hormonal control (in particular via thyroid hormone regulation and insulin signaling) across all male datasets (Figures 3 to 5). Common KEGG pathways across the adult male PCA, LTA, and MTA were limited to inflammatory processes, while the only similarity between the adult female PCA and MTA was the annotation of ‘iron ion import across plasma membrane’ (Figure 6).

**Figure 6.**
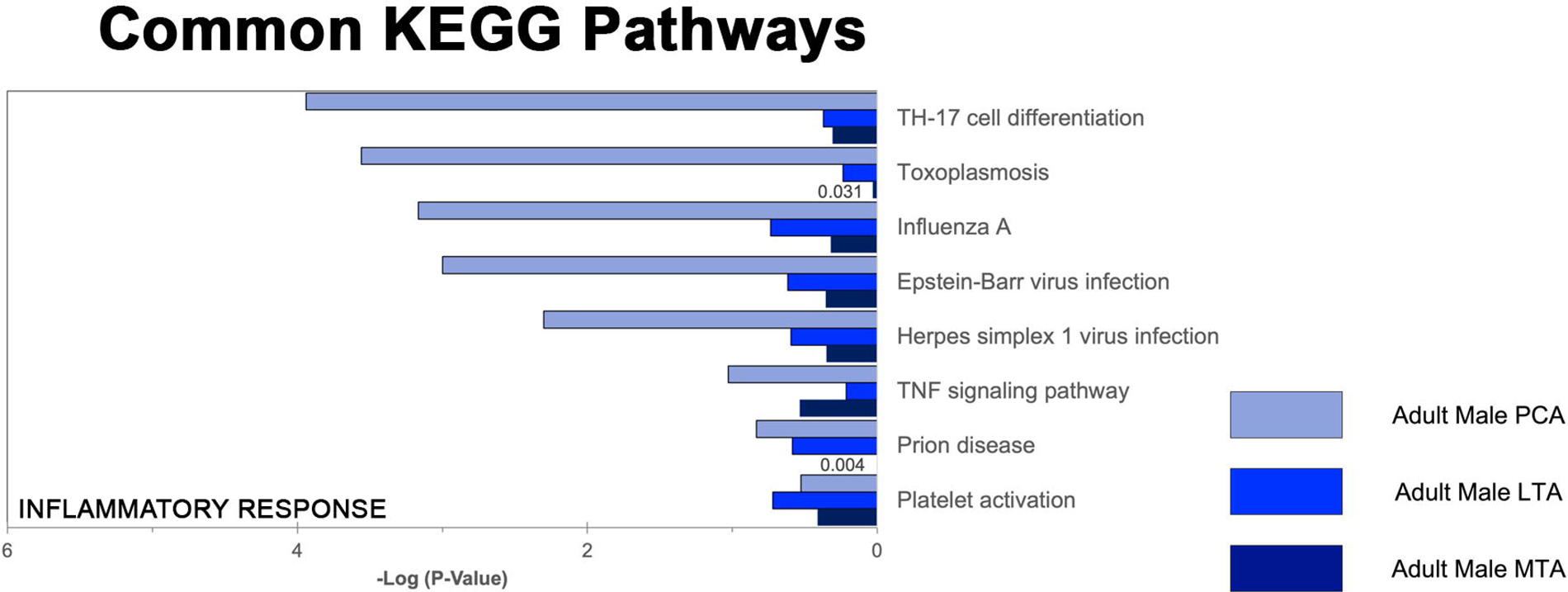
Common KEGG pathways enriched across the PCA, LTA, and MTA muscles. (A) Enriched biological pathways that intersect all laryngeal muscle types were found in the adult male datasets. The adult male PCA, LTA, and MTA datasets only showed an association with pathways involved in the inflammatory response.

## DISCUSSION

The left-right ILM transcriptome profile indicates that laterality is influenced by age, sex, and tissue-specific differences. Reviewing the KEGG Pathways and GO Classifications of DEGs across all experimental conditions allows for identifying genes that may contribute to the intrinsic laterality of laryngeal innervation and reinnervation. The results of this study indicated differences in transcriptional regulation between the left-right abductor (PCA) and adductor (LTA and MTA) muscles. Of the global female and male conditions only 3.0% and 4.8% of left versus right DEGs were found in the abductor muscle compared to the adductor, respectively. Despite having no overlap of DEGs across the left-right PCA, LTA, and MTA muscles, similar regulatory mechanisms between the arytenoid adductors (LTA and MTA) suggest diverse and systematic molecular differences underlying laterality in the healthy state (Figure 4B, 6).

Differential expression analysis demonstrates two general influences regulating the laterality of ILM innervation in uninjured tissue: myogenic and neurophysiological factors. For instance, the most common pathways identified across the global data included ‘axon guidance’ and the ‘calcium signaling pathway’. The GO and KEGG pathways of the implicated DEGs also suggest they exert their influence via inflammatory responses and hormonal control (Figures 3-5). This finding corresponds with the reliance on immune response and hormone signals to promote peripheral nerve regeneration after injury. In the former case, functional recovery of the peripheral nerve tissue is facilitated by inflammatory-related genes in an innate immune response^[30]^. Neurotrophic and guidance factors are upregulated during nerve regeneration, while myelin-associated inhibitors are cleared to promote distal neurite outgrowth to the original targets^[31]^. In the latter event, recent studies have evidenced hormonal control as an essential component of muscle reinnervation after nerve injury. Of relevance, growth hormone treatment in the rat forelimb was associated with significantly attenuated muscle atrophy and enhanced nerve regeneration, muscle reinnervation, and motor function^[32]^. Thyroid hormone and its receptors, localized in Schwann cells, are also critical for promoting nerve regeneration^[33]^. The absence of DEGs related to inflammation in P15 muscles may reflect the development of the immune system between P15 and adulthood^[34]^. Indeed, this phenomenon would agree with age-dependent and sex-related studies concerning musculature hormone control, though further analysis is needed to justify this association^[11, 13, 35-38]^.

Possible limitations of this study are related to the (1) testing of ILM via bulk rather than single-cell RNA-seq, (2) the risk of containing fibers from adjacent muscle or mucosa during microdissection of the laryngeal muscles, and (3) the tissue pooling size. Differences in DEGs with respect to laterality were more strongly detected between the gene sets rather than at the level of the individual genes. Identifying subtle differences in gene expression will require validation of putative targets via functional assays, such as RT-PCR. Inaccurate muscle dissection could conflate the sequencing results with small amounts of adjacent muscle or mucosa contaminating a sample. While confident in our microsurgical techniques, this study recognizes the inherent difficulty in isolating the left from the right PCA at the midline raphe and removing all remnants of mucosa from the left and right MTA. Normalization and downweighing of sample outliers were performed during the quality control step to minimize the impact of this issue. The pooling strategy was implemented to systematically obtain sufficient high-quality RNA required muscle aggregation from three distinct rats. Due to the influence of multiple transcripts per sample, sequencing results could be perturbed by the overrepresentation of one transcriptome profile over another. Pooling bias for RNA can be reduced by increasing the number of biological replicates or increasing the pooled sample size to at least eight tissues to reduce the false discovery rate ^[29, 39, 40]^.

## CONCLUSION

Differential expression analysis of the left-right ILM transcriptome identified age, sex, and muscle-dependent differences in the healthy state. Several common processes implicated in the laterality of ILM gene expression have been shown to influence innervation, These factors include neurophysiological and myogenic processes, inflammatory response, and hormonal control. The genes of interest identified in this study will be evaluated after RLN injury and in models of reinnervation to study their impact on RLN reinnervation. The findings from these investigations may be also critical to developing enhanced treatments to address several neuromuscular degenerative diseases and other muscle-related pathologies.

